# A cross-cultural study of sex-typicality and averageness: Correlation between frontal and lateral measures of human faces

**DOI:** 10.1101/194662

**Authors:** Dariusz P. Danel, Jaroslava Varella Valentova, Oscar R. Sánchez, Juan D. Leongómez, Marco A. C. Varella, Karel Kleisner

**Author notes:** Corresponding author Dariusz Danel.

## Abstract

**Objectives:** Facial averageness and sexual dimorphism are extensively studied attractiveness markers, which are viewed as possible indicators of biological quality. Both are complex morphological traits and both can be easily assessed from frontal and lateral projection of a human face. Interestingly, examination of mutual relations between the frontal and lateral dimensions of these markers has so far received little attention in published research.

**Methods:** In our cross-cultural study, we used geometric morphometric data from male and female faces from Brazil, Cameroon, Colombia, and the Czech Republic, and analysed correlations between frontal and lateral measurements of averageness and degree of maleness/femaleness, i.e. individual variation in features that characterize sexual dimorphism. We also analysed whether the association between frontal and lateral measurements differs in men and women.

**Results:** In general, our results showed a moderate correlation in sexually dimorphic features between lateral and frontal facial configuration in both sexes, while frontal and lateral facial averageness was moderately correlated only in women. This pattern was less consistent when individual populations were analysed separately.

**Conclusions:** Referring to the multiple signalling theory, we propose that especially in women, lateral/frontal correlations support a hypothesis of redundancy of information provided by the frontal and lateral dimension of a given facial attractiveness marker. The absence of a significant correlation in male facial averageness suggests that frontal and lateral averageness may convey different information about an individual.

## 1. INTRODUCTION

Comprehensive research of the role of human face in mate choice led to the identification of several physical markers of facial attractiveness. Two well-known examples of such markers are averageness and sex-typicality (Kościński, 2008; Little, Jones, & DeBruine, 2011; Rhodes, 2006; Thornhill & Gangestad, 1999).

Averageness refers to facial morphology that is biometrically average in a given population (Little et al., 2011; Rhodes, 2006). It has been proposed that average faces are seen as attractive because averageness reflects certain aspects of mate quality, such as heterozygosity, which is in turn related to higher immunocompetence and disease resistance (Gangestad & Buss, 1993; Grammer & Thornhill, 1994; Thornhill & Gangestad, 1993). While the existence of a common genetic component for averageness and attractiveness is still a matter of discussion (Lee et al., 2016; Lie, Rhodes, & Simmons, 2008), the notion of a link between averageness and actual health is supported by some published research (Rhodes et al., 2001; Thornhill & Møller, 1997; Zebrowitz & Rhodes, 2004).

Sex-typicality or, in other words, sexual dimorphism in human faces develops especially during puberty due to the influence of androgens and oestrogens (Enlow, 1990; Farkas, 1988; Tanner, 1989). High levels of testosterone are linked to the development of typically male (i.e., masculine) secondary sexual facial characteristics, such as large jawbones, prominent cheekbones, robust eyebrow ridges, and generally longer and more protruding facial bones. Conversely, the absence of masculinity markers, accompanied by increased size of the lips and thin and receding cheeks, are attributed to a high oestrogen to testosterone ratio and are characteristic of feminine facial morphology (Little et al., 2011; Mydlová, Dupej, Koudelová, & Velemínská, 2015; Rhodes, 2006; Thornhill & Møller, 1997; Verdonck, Gaethofs, Carels, & de Zegher, 1999). Several empirical studies confirm the link between sex-specific facial shapes and steroid hormones in both sexes (Law Smith et al., 2006; Penton-Voak & Chen, 2004; Verdonck et al., 1999). Sexually dimorphic, feminine facial shapes are preferred in female faces because femininity may indicate sexual maturity and a high female reproductive capacity (Gangestad & Scheyd, 2005; Kościński, 2007; Little et al., 2011; Rhodes, 2006). Association between attractiveness and sexual dimorphism in male faces is more complex because in men, both masculine and feminine faces can be preferred (Kościński, 2007; Little et al., 2011; Rhodes, 2006). Several interpretations of aesthetic preference for male sexually dimorphic facial characteristics have been proposed. Firstly, since fully developed sexually dimorphic facial shapes indicate sexual maturity, sexual dimorphism may indicate the ability to produce offspring (Rhodes, 2006; Thornhill & Møller, 1997). Secondly, it has been proposed that highly dimorphic faces advertise immunocompetence: since high levels of male sex hormones are considered detrimental to the immune system, only high-quality individuals can afford to bear the costs of conspicuous sexual ornaments (Folstad & Karter, 1992; Thornhill & Møller, 1997; Wedekind & Folstad, 1994; Zahavi, 1975, but see also Scott, Clark, Boothroyd, & Penton-Voak, 2013 for a discussion of the immunocompetence handicap hypothesis). And finally, exaggerated sexual dimorphism in faces may serve as a cue for personality. For instance, high masculinity in male faces has been associated with higher levels of dominance, aggressiveness, and dishonesty, as well as lower levels of emotionality, warmth, cooperativeness, and parenting abilities (Perrett et al., 1998; Scott et al., 2013). The perception of such anti-social traits may direct women’s preferences towards more feminine male faces (Perrett et al., 1998). Interestingly, preferences for facial sexual dimorphism in both male and female faces may also reflect the recently evolved information-processing mechanisms operating in urbanised and highly populated environments (Scott et al., 2013).

Both facial averageness and secondary sexual characteristics are attractiveness markers that can be easily assessed in facial profiles, but it should be noted that research of facial beauty rarely focuses on lateral facial morphology (Kościński, 2009). Nonetheless, results from several studies of averageness of facial profiles are analogous to the results of en-face investigations and they do show that mathematically average profiles are attractive (Spyropoulos & Halazonetis, 2001; Valentine, Darling, & Donnelly, 2004; Valenzano, Mennucci, Tartarelli, & Cellerino, 2006). Moreover, numerous studies of facial profiles carried out within aesthetic medicine and orthodontics found that morphological deviations from average profile shapes (i.e. orthognathic profiles with biometrically normal proportions) lower the perceived facial attractiveness (e.g. Gautam, Shashikalakumari, & Garg, 2013; Hönn, Dietz, Eiselt, & Göz, 2008; Johnston et al., 2005; Soh, Chew, & Wong, 2007; Türkkahraman & Gökalp, 2004). Similarly, research results on the attractiveness of facial profile sexual dimorphism are also in line with some findings of studies of en-face facial morphology. In particular, higher femininity of both male and female facial profiles was found to be associated with higher facial attractiveness (Swaddle & Reierson, 2002; Valenzano et al., 2006) and, at least in men, also with lower perceived dominance (Swaddle & Reierson, 2002).

The agreement between results obtained by en-face and facial profile studies implies that lateral and frontal measures of averageness and sexual dimorphism should be correlated. Such a view corresponds with anatomical and developmental analyses which show that horizontal, vertical, and anteroposterior craniofacial dimensions are interrelated (Enlow, 1990). On the other hand, faces are complex and multi-trait body parts that cannot be studied as one developmentally homogenous structure. Indeed, the morphogenesis of particular facial characteristics is subjected to genetic, epigenetic, environmental, and even functional influences (Claes et al., 2014; Enlow, 1990; Hallgrimsson, Mio, Marcucio, & Spritz, 2014; Liu et al., 2012). Multilevel interactions between these factors result in a considerable morphological diversity of particular facial features and faces as a whole (Enlow, 1990; Hallgrimsson et al., 2014). At least in theory, therefore, measurements of seemingly identical qualities from frontal and lateral perspective could be uncorrelated or correlate only weakly. In fact, a three-dimensional computer tomography study of human skulls had shown that mutual relations between the size of different facial structures can be complex, and a straightforward anticipation of correlation between traits dimensions is not necessarily justified (Wang, Otsuka, Akimoto, & Sato, 2013).

To our best knowledge, only one study analysed directly the association between frontal and lateral markers of facial attractiveness, such as averageness and sexual dimorphism. Using frontal and lateral facial photographs of the same individual and applying geometric morphometric to analyse facial morphology, Danel, Dziedzic-Danel, and Kleisner (2016) found that neither individual expression of frontal averageness nor frontal sexually dimorphic traits are correlated with their lateral counterparts. The absence of statistically significant correlations between frontal and lateral measurements was found in both male and female faces. However, the sample of facial pictures used by Danel et al. (2016) was relatively small and limited only to one population, namely adult Caucasians of Polish origin. These shortcomings, jointly with the absence of other studies on interrelations between frontal and lateral markers of facial attractiveness, do not allow for a generalization the findings of Danel et al. (2016) or an extrapolation of this study’s conclusions to different populations.

In the current study, we address the abovementioned limitations and replicate Danel’s et al. (2016) study using a larger and more ethnically diverse sample, which includes participants from the Czech Republic (Europe), Cameroon (Africa), and two South American countries: Brazil and Colombia. More specifically, we analyse correlations between frontal and lateral measures of averageness and individual expression of sexually dimorphic traits in these four different populations from different parts of the world. We started by testing whether frontal and lateral views provide redundant cues about an individual’s sex-typicality and averageness. If so, the individual expression of averageness and level of sexual dimorphism (maleness/ femaleness) should be closely correlated. Then we examined whether the association between frontal and lateral measurements differs in male and female faces.

## 2. METHODS

### 2.1 Ethics statement

These particular studies were approved by the Institutional Review Board of Charles University, Faculty of Science (for the data collection in Brazil – 2011/7, for the data collection in Cameroon and the Czech Republic – 2014/10) and the Institutional Committee on Research Ethics, El Bosque University (for the data collection in Colombia – 2016/2). All participants provided a written informed consent prior to participating in the study. Morphometric data of frontal and lateral images were analysed anonymously.

### 2.2 Facial photographs

We used facial frontal and lateral portraits of 100 Brazilians (51 women: Mean Age±SD = 24.1±4.9, range 18-35; 49 men: Mean Age±SD = 23.29±3.63, range 18-32), 102 Cameroonians (51 women: Mean Age±SD = 24.92±8.4, range: 17-54; 52 men: Mean Age±SD = 23.75±5.49, range 17-44), 98 Colombians (49 women: Mean Age±SD = 21.5±3.29, range 18-33; 49 men: Mean Age±SD = 20.1± 1.71, range 18-27), and 191 Czechs (139 women: Mean Age±SD = 21.45±2.51, range 18-34; 52 men: Mean Age±SD = 23.06±4.45, range 19-39). Due to a possible laterality of human facial morphology (Danel & Pawlowski, 2007; Farkas & Cheung, 1981; Simmons, Rhodes, Peters, & Koehler, 2004), we decided to use always the same, namely left, facial profile. Although facial photographs were collected by different authors, they followed a standardised procedure (Trebicky, Fialová, Kleisner, & Havlícek, 2016). As stated above, our dataset is highly ethnically diverse, since it includes individuals from one European, one African, and two ethnically mixed South American countries.

### 2.3 Geometric morphometries

We determined 72 landmarks (including 36 semilandmarks) on frontal facial portraits and and 21 landmarks (including 6 semilandmarks) on profile images. Landmarks are corresponding locations that can be anatomically or geometrically defined in all objects in the sample. While landmarks reflect homologous structures and locations on faces of different individuals, semilandmarks denote curves and outlines. For landmark and semilandmarks locations on frontal and lateral facial images, see Danel et al. (2016).

Facial frontal and profile configurations of landmarks were superimposed by a generalised Procrustes analysis (GPA) using the ‘gpagen’ function implemented in the geomorph package in R (Adams & Otárola-Castillo, 2013). GPA standardized the size of the objects and removed rotational and translational effects in order to minimize distances between homologous landmarks. The ‘gpagen’ function was also used to align sliding semilandmarks by means on the minimum bending energy criterion.

For each set of faces we run separate GPA and computed the mean configuration (consensus). The Procrustes distances between the consesus and each configuration in the set was computed and used as a measure of individual averageness A higher facial averageness score thus indicates closer proximity of a configuration to the mean shape.

To assess the level of individual expression of facial traits responsible for sexual shape dimorphism (SShD), i.e., the degree of maleness/femaleness as suggested by Mitteorecker et al. (2015), we determined the position of an individual facial shape along the axis between male and female mean shapes (Mitteroecker et al., 2015; Valenzano et al., 2006). This position can be numerically expressed by continuous scores of ordination constrained by sex, i.e., by projecting individual facial configurations onto the vector between the average female and male face.

### 2.4 Statistical analysis

Since the individual scores of facial metrics did not meet the statistical requirement for normal distribution, we performed correlational analyses using nonparametric methods. Relationships between frontal and lateral facial configurations were investigated using Kendall’s Tau correlation coefficient for measurements of both averageness and sexual shape dimorphism. Comparisons of correlation coefficients obtained for a pooled sample of men and women were carried out using Fisher’s transformation for Kendall’s Tau coefficients according to Walker (2003) and Zar (1999). All these tests are two-tailed.

## 3. RESULTS

Correlations between frontal and lateral facial shapes for averageness and individual expression of sexually dimorphic traits (maleness/femaleness) are summarised in Table 1. When shape coordinates for all samples were pooled together across countries but separately for men and women, averageness showed a significant frontal/lateral association only in women (Fig 1). A formal comparison of sex differences in the correlation coefficients confirmed that the association between frontal and lateral facial averageness is significantly stronger in women than in men (Z = 4.11, p < .0001).

**Table 1.** Kendall’s Tau correlations between frontal and lateral facial configuration for averageness (AVRGN) and the degree individual expression of traits associated with sexual shape dimorphism (SShD).

| SAMPLE | SEX | AVRGN | SShD | N |
| --- | --- | --- | --- | --- |
| ALL | women | .275** | .200** | 290 |
|  | men | .042 | .202** | 201 |
| Brazil | women | .126 | .194* | 51 |
|  | men | .064 | .092 | 49 |
| Cameroon | women | .220* | .154 | 51 |
|  | men | -.035 | .071 | 51 |
| Colombia | women | .143 | .197* | 49 |
|  | men | .022 | .224* | 49 |
| Czech Republic | women | .206** | .260** | 139 |
|  | men | .015 | .101 | 52 |

\* p-value $\leq .05$ \*\* p-value $\leq .001$

On the other hand, the association between frontal and lateral individual expression of sexual shape dimorphism was significant in the pooled samples of both men and women (Fig 1). The strength of the correlation was comparable for both women and men, and difference between the correlations was not statistically significant (Z = −0.04, p = .971).

**Figure 1.**
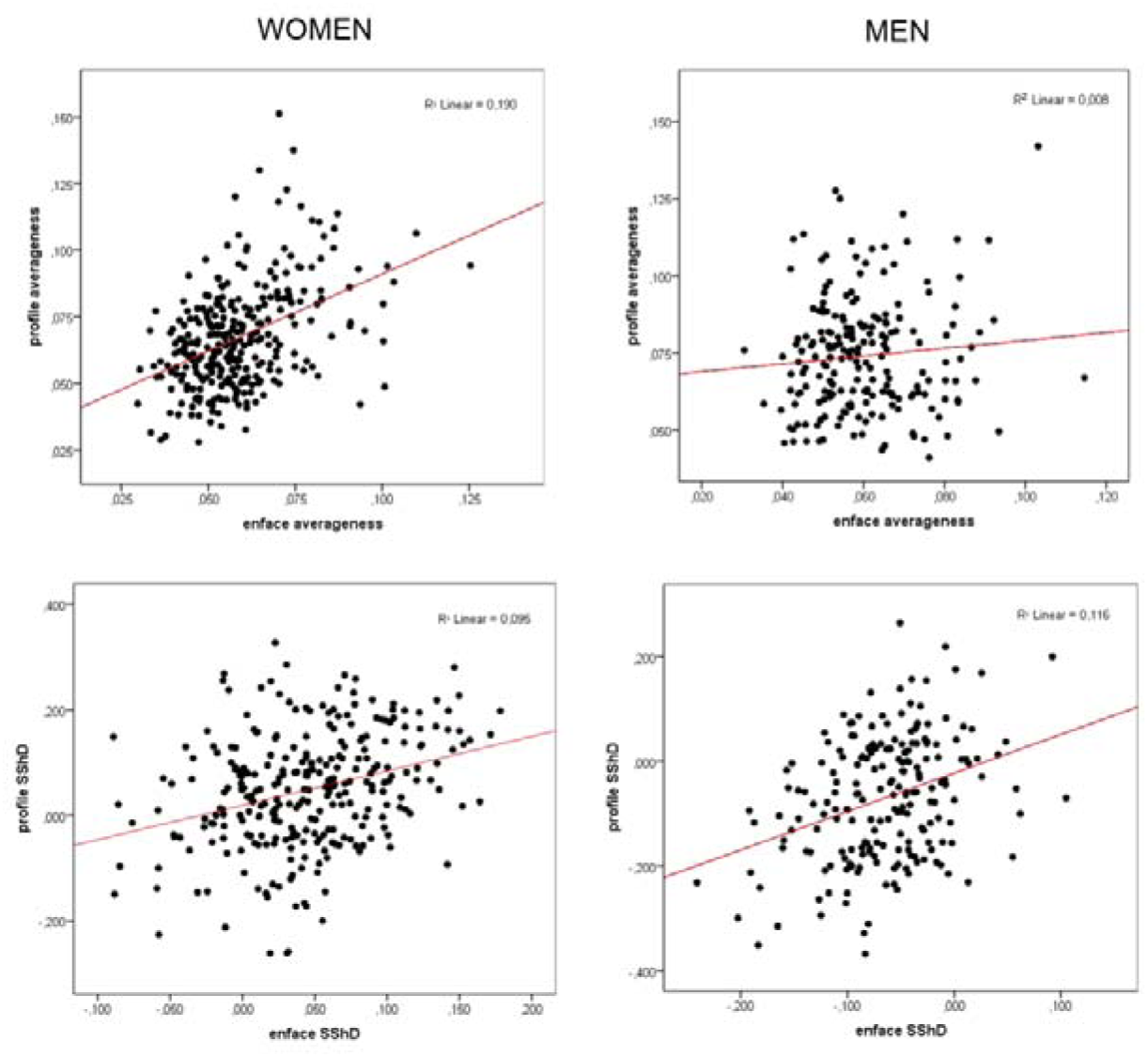
Scatterplots showing pooled data separately for men and women. Upper panel shows correlations between frontal and lateral values of averageness (i.e., the distance of each specimen from the Procrustes mean configuration). Lower panel demonstrates the association between individual scores of sexual shape dimorphism (SShD) computed for frontal and lateral configurations.

When samples were analysed separately, it turned out that Brazilian women display a significant frontal-lateral correlation for femaleness but not for averageness. Conversely, analysis of the sample of Cameroonian women showed a significant frontal-lateral correlation for averageness but not for femaleness. In the Colombian women, neither frontal-lateral averageness nor measures of femaleness were significantly correlated. In Czech women, frontal configurations were significantly correlated with profiles in both averageness and femaleness.

In the male samples, the correlation coefficient for frontal and lateral maleness reached significant levels only in the Colombian men. In the other populations, the association between frontal and lateral facial configurations was not statistically significant.

In general, female faces showed a higher degree of correlation between frontal and lateral configuration for both averageness and individual expression of sexually dimorphic shape features than men. An exception was the Colombian sample, where correlation coefficients for measures of sexual shape dimorphism was higher for men than for women.

## 4. DICUSSION

Using data from four populations from different parts of the world, we examined whether profile and lateral configurations correlate in individual expression of averageness and sexually dimorphic characteristics of facial shape (maleness and femaleness). In general, we found that individual expression of profile and frontal sexual dimorphism is moderately correlated in both men and women. The strength of these associations does not differ between men and women. Correlations between frontal and lateral averageness did, however, reveal some profound differences between the sexes: while in women, averageness of enfaces was significantly associated with averageness of profiles, no such correlation was found in men.

Analyses performed separately for particular populations showed a similar trend but the association was significant only in some cases: while frontal and lateral averageness were significantly correlated in Czech and Cameroonian women (and not related in other groups), frontal and lateral measures of sexual shape dimorphism were significantly associated in Czech and Brazilian women and in Colombian men. The observed absence of homogenous pattern in particular samples might be due to the relatively small sample sizes of the particular analysed populations.

The current results, which show significant correlations between frontal and lateral facial configurations, are in line with ontogenetic analyses indicating that various facial structures are developmentally interrelated (Enlow, 1990). Our results are also in agreement with facial attractiveness studies which show that frontal and lateral facial averageness and sexual dimorphism may exert congruent effects on the facial beauty of individuals (Swaddle & Reierson, 2002; Valentine et al., 2004; Valenzano et al., 2006).

Nonetheless, it should be noted that the correlations found in our study are at most moderate. This indicates that the spatial development of complex facial configurations, such as frontal and lateral facial averageness (e.g. Claes & Shriver, 2014; Enlow, 1990; Hallgrimsson et al., 2014; Liu et al., 2012), may be influenced by various factors. A possible explanation of the only moderate correlations between lateral and frontal facial structures might be that faces show a higher phenotypic variation and lower between-trait correlation that other parts of the human body. It has been proposed that this morphological diversity could be the consequence of human sociality and selection pressures on identity signalling facilitated by facial recognition in complex social systems (Sheehan et al., 2014).

However, even if individual recognition and discrimination does play a role in the development of highly variable and loosely related frontal and lateral facial morphology, it does not explain why these correlations are generally weaker in men than in women. We propose that these sex differences between women and men may reflect differences in the information conveyed by male and female faces. Previous studies of multimodal signalling show that vocal and facial attractiveness are often correlated in women (Feinberg et al., 2005; Penton-Voak et al., 2001), while in men the evidence is more ambiguous (Lander, 2008; Oguchi & Kikuchi, 1997; Saxton, Burriss, Murray, Rowland, & Roberts, 2009; Saxton, Caryl, & Roberts, 2006; Valentová, Roberts, & Havlícek, 2013; Wells, Baguley, Sergeant, & Dunn, 2013). Faces are not multimodal but unimodal entities which depend solely on visual perception. Nevertheless, even unimodal objects, such as faces, can still carry multiple signals which can be either redundant or non-redundant. While redundant signals bear identical information, in non-redundant signals, different components covey different information (Johnstone & Grafen, 1992; Partan & Marler, 2005; Schluter & Price, 1993). A stronger correlation between frontal and lateral structures should point to signal redundancy, while weak or no correlation may indicate a disparity of information carried by the relevant features. According to our results regarding averageness, it seems that female faces may carry more redundant signals since in women, lateral and frontal structures are more closely correlated. Male faces, on the other hand, show no correlation between frontal and lateral averageness, which may indicate that the two aspects of the overall facial morphology convey different messages.

Differences between the sexes in frontal-lateral correlation in averageness can be explained by the fact that female faces are under a strong intersexual and intrasexual selection pressure for physical attractiveness (Buss et al., 1989; Fisher, 2004; Hatfield & Sprecher, 1995; Li, Bailey, Kenrick, & Linsenmeier, 2002). Because faces closer to the average are commonly seen as more attractive (Little et al., 2011), information provided by women’s frontal facial structures may be further reinforced if the profile morphology is also closer to an average shape. In men, however, selection pressure on their facial morphology is weaker, because other modalities – such as voice pitch, social status, career prospects, and access to resources – determine the overall mate value of the individual and affect women’s preferences (Buss et al., 1989; Feingold, 1992; Hatfield & Sprecher, 1995; Li et al., 2002; Saxton, Mackey, McCarty, & Neave, 2016). A lower correlation between frontal and lateral measures of facial averageness in men may imply that the two facets of averageness carry non-redundant information (possibly due to lower selection pressure on averageness of male faces). In the case of sexual shape dimorphism, frontal morphology correlates with lateral in both sexes, which suggests a redundancy of information conveyed by profile and lateral facial configurations. This could be because sexual dimorphism is an important facial cue in both sexes, and facial features associated with sex-typicality are under intersexual and/or intrasexual selection pressure in both men and women (Fink, Klappauf, Brewer, & Shackelford, 2014; Puts, 2010; Scott et al., 2013).

Multiple signalling theory, which predicts that depending on the redundant and non-redundant character of a signal the same or different information can be conveyed by different features (Wells, Dunn, Sergeant, & Davies, 2009), offers an appropriate framework to interpret the presence or absence of correlations observed in the current study. Our study did not, however, test any explicit hypotheses which could be derived from this general theory. Future research could address this issue by, for instance, investigating whether the qualities of individuals predicted based on frontal images are in agreement with analogous assessments made on the basis of facial profiles. As in the current examination, implementation of such studies in cross-cultural settings may also provide a broader and more universal perspective on factors which might influence the appearance and perception of various facial cues.

As for limitations, our sample sizes for men and women in particular populations were relatively small. Simple sample size estimations show that an adequate sample size for identification of a significant association of Kendall’s Tau = .15 (equivalent of r = .23; Gilpin, 1993; Walker, 2003), with statistical power of 1 − β = .90, should be over n = 190 (Hulley, Cummings, Browner, Grady, & Newman, 2013). Such numbers of men and women were obtained when data from the four populations were pooled together, which also implies that results from the separate populations should be interpreted with a degree of caution. Furthermore, in this study only left profile pictures were taken. Faces are not, however, perfectly bilateral (Danel & Pawlowski, 2007; Farkas & Cheung, 1981; Simmons et al., 2004), and analyses of correlations between frontal and lateral faces should therefore be replicated also with right profiles.

In conclusion, the current cross-cultural study examined interrelations between frontal and lateral configurations of averageness and sexual shape dimorphism, two commonly accepted markers of facial attractiveness. Results show that measures of frontal and lateral shape dimorphism are moderately correlated in both men and women. A similar moderate correlation was observed in frontal and lateral facial averageness in women but not in men. The framework provided by multiple signalling theory may help explain the observed interrelations, but further research focused specifically on the redundant and non-redundant signalling roles of traits associated with lateral and frontal averageness and sexual shape dimorphism is needed.

## ACKNOWLEDGEMENTS

We would like to thank to Tomáš Kočnar, Robert Mbe Akkoko, and Šimon Pokorný for their help with collection of Cameroonian and Czech data. Our thanks belong also to Petr Tureček for his advices on statistical data analyses. The authors have no conflict of interest to declare.

## AUTHOR CONTRIBUTION

DD and KK analyzed the data and drafted the manuscript. DD and KK designed the study, and directed implementation and data collection. JVL, ORS, JDL, MACV and KK collected the data. JVL, ORS, JDL, and MACV edited the manuscript for intellectual content and provided critical comments on the manuscript.

